# Selective modulation of population dynamics during neuroprosthetic skill learning

**DOI:** 10.1101/2021.01.08.425917

**Authors:** Ellen L. Zippi, Albert K. You, Karunesh Ganguly, Jose M. Carmena

**Affiliations:** Helen Wills Neuroscience Institute, University of California Berkeley, Berkeley, California, USA; Department of Electrical Engineering and Computer Sciences, University of California Berkeley, Berkeley, California, USA; Neurology & Rehabilitation Service, San Francisco VA Medical Center, San Francisco, CA, USA; Department of Neurology, University of California, San Francisco, CA, USA

## Abstract

Learning to control a brain-machine interface (BMI) is associated with the emergence of coordinated neural dynamics in populations of neurons whose activity serves as direct input to the BMI decoder (direct subpopulation). While previous work shows differential modification of firing rate modulation in this population relative to a population whose activity was not directly input to the BMI decoder (indirect subpopulation), little is known about how learning-rated changes in cortical population dynamics within these groups compare. To investigate this, we monitored both direct and indirect subpopulations as two macaque monkeys learned to control a BMI. We found that while the combined population increased coordinated neural dynamics, this coordination was primarily driven by changes in the direct subpopulation while the indirect subpopulation remained relatively stable. These findings indicate that motor cortex refines cortical dynamics throughout the entire network during learning, with a more pronounced effect in ensembles causally linked to behavior.

## Introduction

Refinement of learned behaviors depends on mechanisms of reinforcement learning involving many cortical and subcortical structures (Donchin et al., 2012; Krakauer et al., 2004; Sing & Smith, 2010; Sutton & Barto, 1998). Just as specific behavioral actions can be reinforced, the specific cortical population activity required to produce these actions efficiently and with less variability can also be reinforced (Athalye et al., 2018; Costa, 2011; Tumer & Brainard, 2007). The coordinated activity of this cortical population is often characterized by low-dimensional dynamics (Ames & Churchland, 2019; Athalye et al., 2017; Churchland et al., 2012; Golub et al., 2018; Heming et al., 2019; Kao et al., 2015; Oby et al., 2019; Sadtler et al., 2014). These dynamics capture the patterns of co-activation across neurons within the population driving behavior. Changes in these cortical dynamics have been linked to learning novel actions.

It has been proposed that two parallel mechanisms of learning act to reinforce specific cortical population dynamics; fast reinforcement of the cortical dynamics that naturally produce a desired behavior and slower reinforcement that refines them to result in more reliable production of neural activity patterns driving the desired behavior (Athalye et al., 2020; Dayan & Cohen, 2011). Studying the mechanisms of cortical reinforcement that underlie behavioral reinforcement can be challenging as the exact neural population controlling the desired behavior, and thus the impact of specific changes in neural firing patterns on that behavior, is unknown. Brain-machine interfaces (BMIs), however, allow the experimenter to precisely define the mapping between recorded neural activity and behavior (Carmena et al., 2003; Shenoy & Carmena, 2014; Taylor, 2002). Depending on the chosen mapping, learning to control a BMI can require an animal to learn novel, cortical dynamics to obtain rapid and precise control (Churchland et al., 2012; Ganguly et al., 2011; Ganguly & Carmena, 2009; Orsborn et al., 2014). Thus, studying how cortical population dynamics change as an animal learns to control a BMI can shed light onto how the dynamics underlying selected novel behaviors are reinforced.

The coordination of activity between neurons can be represented by their covariance structure, which explains how neurons interact within a population (Ames & Churchland, 2019; Athalye et al., 2017; Heming et al., 2019; You et al., 2019). Previous studies have shown neural populations are constrained to generate activity patterns within their pre-existing covariance structure within short timescales (Golub et al., 2018; Sadtler et al., 2014). These results suggest that it is faster to learn to control pre-existing cortical population dynamics than it is to modify them to be outside of the pre-existing covariance structure. Other work has also shown that animals can learn to control a BMI with a decoder that requires neural patterns outside of the pre-existing covariance structure over the course of multiple days (Athalye et al., 2017; Oby et al., 2019). Learning to control these decoders involved an increase in coordinated activity and decrease in neural variability. The modification of the cortical dynamics required for efficient control requires a longer timescale than the fast reassociation of cortical population dynamics. This suggests that learning new skills involves the production of new underlying cortical population activity and the development of these new population dynamics occurs over a longer timescale. As the dynamics that produce desirable outcomes evolve, they are reinforced and associated with the outcome.

When a motor cortical BMI is designed, the experimenter selects only a small subset of neurons from motor cortex to use as input to the decoder (direct neurons). These neurons exist within a large network of other motor cortical neurons (indirect neurons). While the selection of direct neurons is arbitrary and there is initially no functional difference between the two groups, previous work has shown that differences in neural activity between direct and indirect neurons emerge with learning. For example, it has been demonstrated that the task-relevant modulation of indirect neurons gradually reduces relative to direct neurons as monkeys learn to control a BMI (Ganguly et al., 2011). Additionally, it has been shown in rodents that coherence develops between dorsal striatum and direct neurons, but not indirect neurons, as the animals improve control (Koralek et al., 2012, 2013; Neely et al., 2018). In a separate study using 2-photon calcium imaging to record neural activity, it was shown that mice modulate activity of direct and indirect neurons in early learning, but eventually learn to mostly modulate activity of the direct neurons (Clancy et al., 2014). Additional primate BMI studies investigating neural adaptation to changes in the decoder found that even within the direct neurons there were differences in the degree of adaptation related to the degree to which the neurons drove behavioral error (Chase et al., 2012; Jarosiewicz et al., 2008; Orsborn et al., 2014). Thus, it is likely that the initial cortical dynamics that produce desirable outcomes involve both the neurons that were selected to control the BMI as well as the surrounding cortical network. Over time, as these cortical dynamics are refined, they may adapt to involve smaller populations and rely less on those that do not drive behavior.

If this hypothesis is true, we might expect differences in the changes in cortical dynamics that occur within direct and indirect neurons. As cortical dynamics are modified to more effectively and efficiently control a BMI, direct neuron population activity would be expected to undergo further modification than indirect neuron population activity as additional modifications to the indirect neuron activity would not directly result in behavioral changes. Here, we investigate this idea by studying recorded ensembles of motor cortical neurons while only a subset was assigned to have a causal role during control of a BMI, and characterize the differential changes in coordinated neural dynamics between the direct and indirect subpopulations.

## Results

Two rhesus macaques (P and R) were chronically implanted with bilateral microelectrode arrays in primary motor and dorsal premotor cortices, with electrodes from a single hemisphere used for BMI control and subsequent analyses (see Methods). The monkeys learned to perform a two-dimensional, self-initiated center-out BMI task, in which they drove a cursor under neural control to one of eight randomly instructed peripheral targets for a juice reward (Figure 1a). The next trial was initiated by driving the cursor back to the center target. Trials from all days of the experiment were concatenated then separated into 150-trial epochs since the number of successful trials was lower in early days of learning. To capture the correlates of learning before performance saturated, only the first 15 epochs were used for each animal.

**Figure 1.**
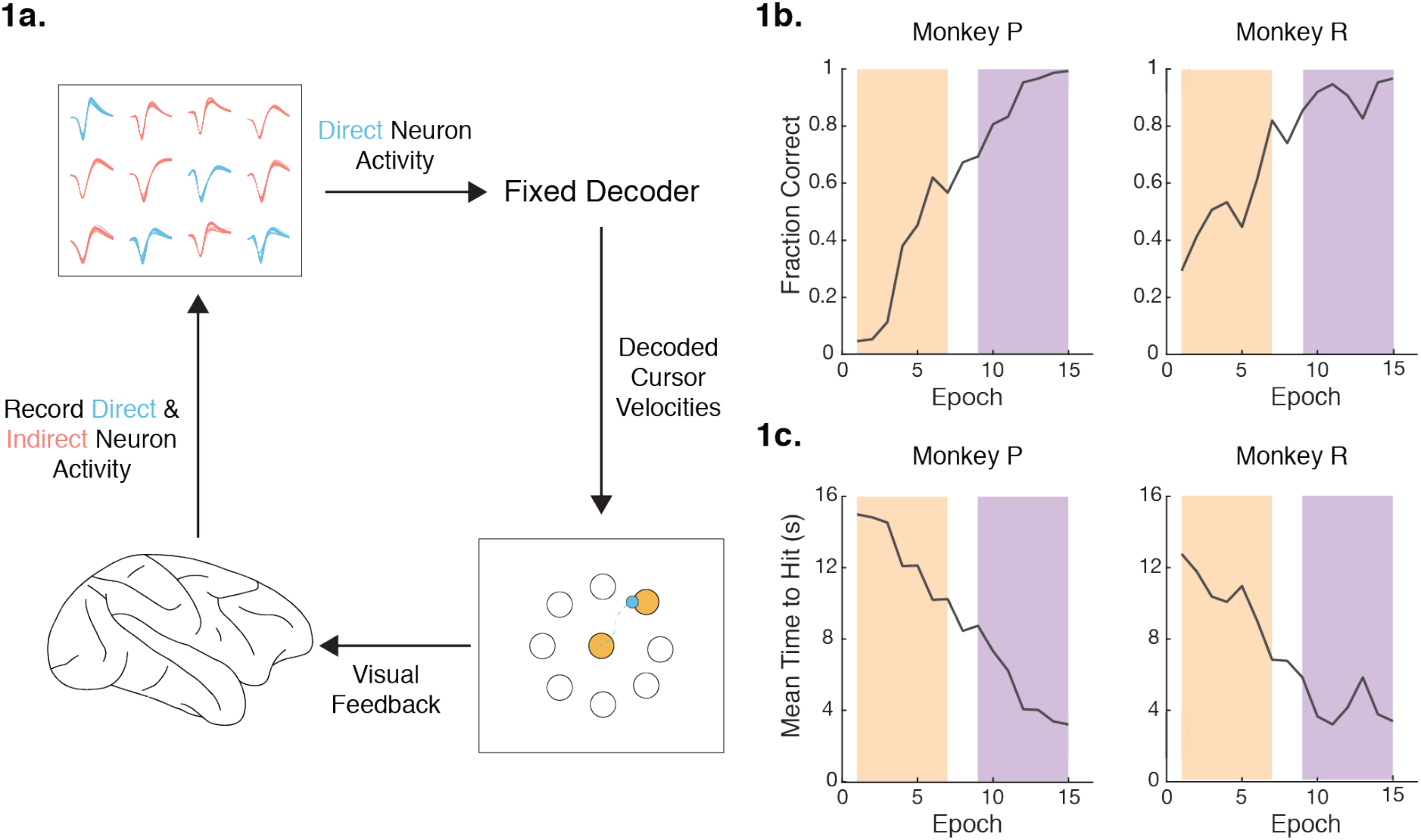
Experimental setup and behavioral performance. (a) Experimental setup from Ganguly & Carmena, 2009. Activity recorded from direct neurons (blue) in M1/PMd was input into a fixed linear decoder and used to drive a computer cursor to perform a center-out task (see Methods for details). Activity from indirect neurons (red) was simultaneously recorded, but was not input into the decoder. (b) Performance improves over the first 15 training epochs for Monkey P and Monkey R. Each training epoch consists of 150 initiated trials. For some analyses, epochs were divided into groups of early (orange) and late (purple). Fraction of initiated trials that were successful increased over training epochs. (c) The time to reach a target decreased over training epochs.

To characterize the neural dynamics associated with learning to control a BMI, we examined recorded ensembles of motor cortical neurons from which only a subset was assigned a causal role during control. We define these neurons, whose activity was used as a direct input to the BMI decoder, as “direct neurons” (Monkey P, N = 15; Monkey R, N = 10). The remaining recorded motor cortical neurons, recorded using the same two 4×4 mm 64-channel microelectrode arrays (interelectrode distance 500 um), whose activity was not used as direct input to the BMI, we define as “indirect neurons” (Monkey P, N = 29-69; Monkey R, N = 87-187). For some analyses in which it is important to consider the same population across epochs we refer to indirect neurons that were stably recorded across all 15 epochs as “stable indirect neurons” (Monkey P, N = 17; Monkey R, N = 14). Stability of the indirect neurons was assessed using the methods described in Fraser & Schwartz, 2012.

First, we examined how the neural firing rate variance changed in each subpopulation over learning. Changes in neural variance are often used as a proxy for neural exploration, as increasing the variance in firing rate allows for neurons to form different coordinated patterns of firing (Athalye et al., 2017). Past work has shown neural activity to fire in coordinated efforts, thereby decreasing the dimensionality in neural space. We commonly refer to these low-dimensional spaces as manifolds or neural subspaces (Athalye et al., 2017; Golub et al., 2018; Oby et al., 2019; Sadtler et al., 2014). Intuitively, increasing the variance of a neuron’s firing rate moves the neuron along or away from the neural subspace, respectively. Epochs were separated between early and late for each animal (Epochs 1-7 and Epochs 9-15, respectively) to track differences as behavioral performance improved. The firing rate for each neuron was binned in 100 ms intervals. The binned firing rate variance for each neuron was then averaged across all neurons for each epoch. Since the number of direct and indirect units differed with varying numbers of indirect neurons recorded each epoch, we normalized the firing rate variance for each subpopulation based on the mean variance in early learning to assess relative change in variance. In both animals, we observed an increase in relative unit variance between early and late learning for both direct and indirect subpopulations (Figure 2). This increase in variance suggests a concerted effort of neural exploration that exists in a broader network that includes both direct and indirect neurons.

**Figure 2.**
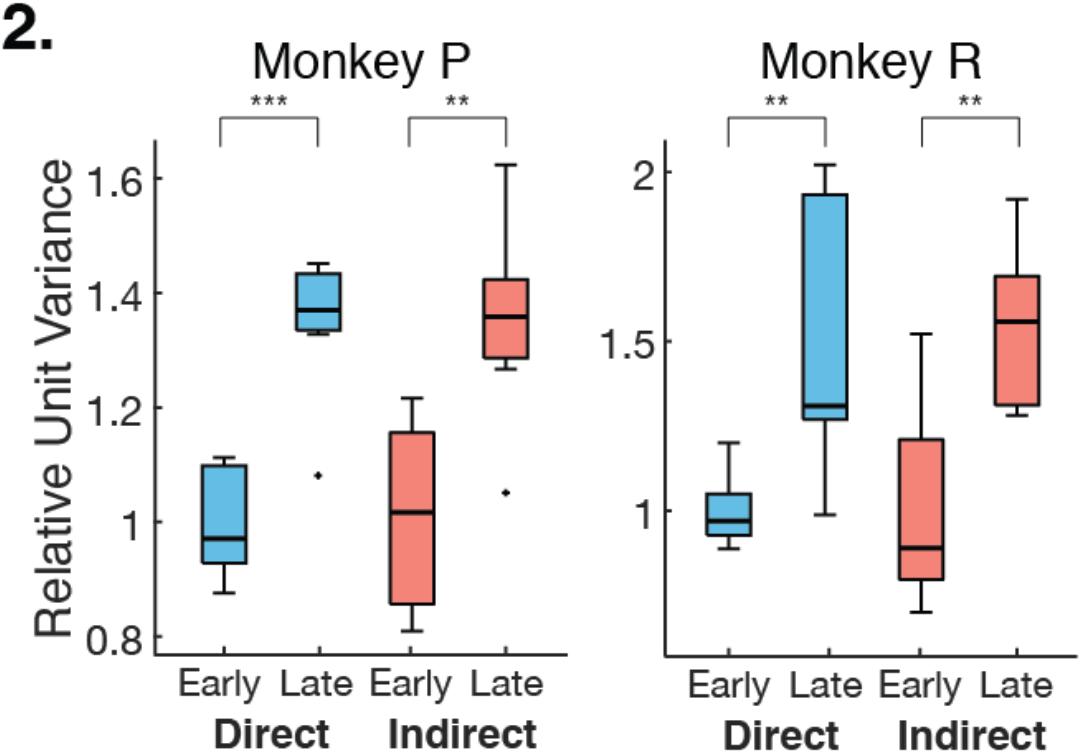
Neural variance increases with learning. Variance was calculated for each neuron and then averaged across neurons. Relative variance was calculated by normalizing to the mean early variance within subpopulation. Both direct and indirect subpopulations increased relative neural variance from early to late learning. (Unpaired t-test; Monkey P: direct p = 8.63e-5, indirect p = 0.002; Monkey R: direct p = 0.007, indirect p = 0.003).

Previous work has shown neurons to fire in increasingly coordinated patterns as performance improves (Athalye et al., 2017). We consider these changes as a proxy for consolidation since the neural variance is consolidating onto low-dimensional subspaces. We use factor analysis (FA) to separate the neural variance in the population into two components - private and shared variances. The shared variance is the variance between neurons in the population and can be thought of as the underlying correlated firing pattern in the recorded population. Conversely, the private variance denotes the amount of variance each neuron has that is independent from the rest of the population. Past studies have explored the roles of these components in the direct neuron population, showing private variance as a proxy for exploration while an increase in shared variance is correlated to skill consolidation. Here we characterized the amount of consolidation as the proportion of the total neural variance that is captured in shared spaces for the combined direct and indirect population. That is, the ratio of shared over total (SOT) variance. Since the indirect population consisted of different units each epoch, we normalized the SOT to the mean SOT for early epochs (Figure 3a). In both animals, we found that the relative SOT increased between early and late learning for the entire recorded population including both direct and indirect neurons. Together, with the increase in variance over learning, our results indicated a high level of consolidation that occurs in the entire recorded population driven by increased exploration as BMI performance improves. In order to assess that the effect was not due to day-to-day differences in the population, we conducted the same analysis on neurons that were stably recorded across all 15 epochs, which yielded consistent results (Figure 3b).

**Figure 3.**
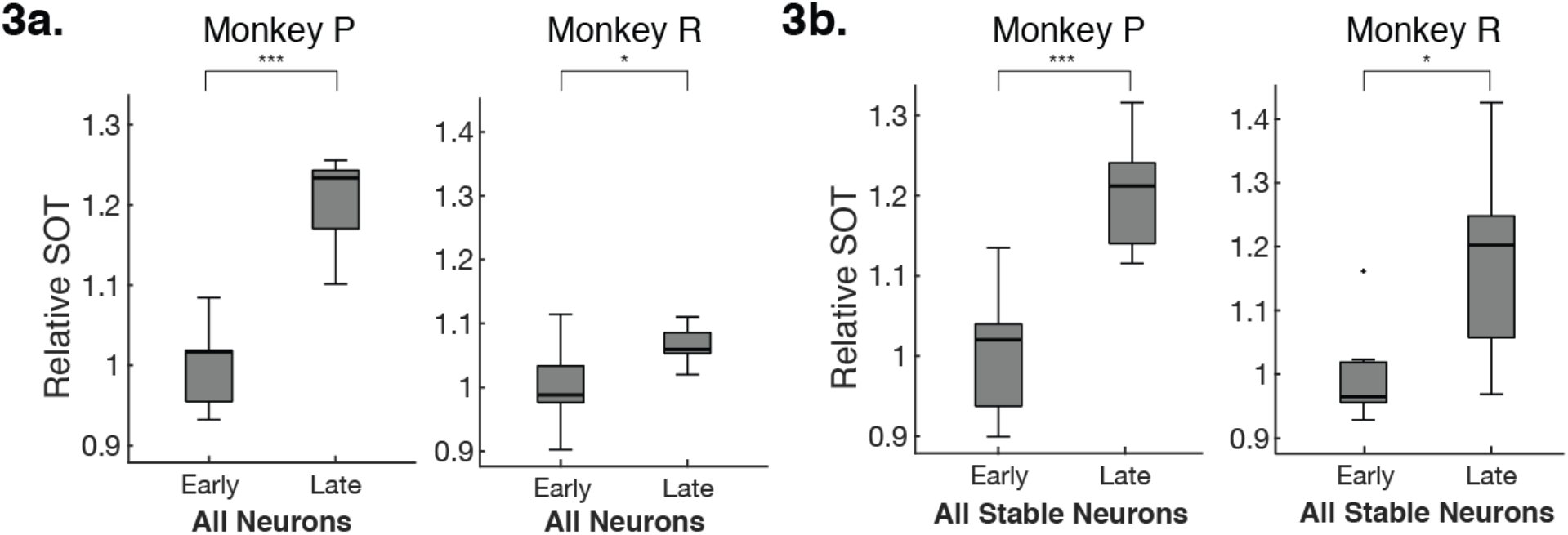
Neural activity consolidates with learning. The relative shared-over-total variance (SOT) ratio was calculated with respect to the mean early SOT within each subpopulation. (a) Relative SOT increases between early and late learning, indicating an overall consolidation of neural activity in the entire recorded population (Unpaired t-test; Monkey P, p = 1.31e-5; Monkey R, p = 0.033). (b) Relative SOT also increases between early and late learning for a stably recorded population consisting of the same units each epoch (Unpaired t-test; Monkey P, p = 2.80e-4; Monkey R, p = 0.020).

While the increase in SOT shows an increase in consolidation in the combined population, it does not explain whether these changes are driven by a specific subpopulation. To answer this question, we considered the partial shared-over-total variance (pSOT) ratio of the stably recorded population to see how the same population of neurons consolidated over learning (see Methods). Intuitively, the pSOT ratio asks how much of the overall neural consolidation was driven by one subpopulation versus the other. We see that, relative to early learning, there was a significant increase in pSOT_direct_ in late learning but not in pSOT_indirect_ (Figure 4), indicating that the increase in coordination of population activity seen across the stably recorded population was driven by the increase in coordination of population activity within the direct subpopulation. This suggests that the increased neural exploration in the network was primarily a consequence of changes in coordinated patterns specific to the direct neurons.

**Figure 4.**
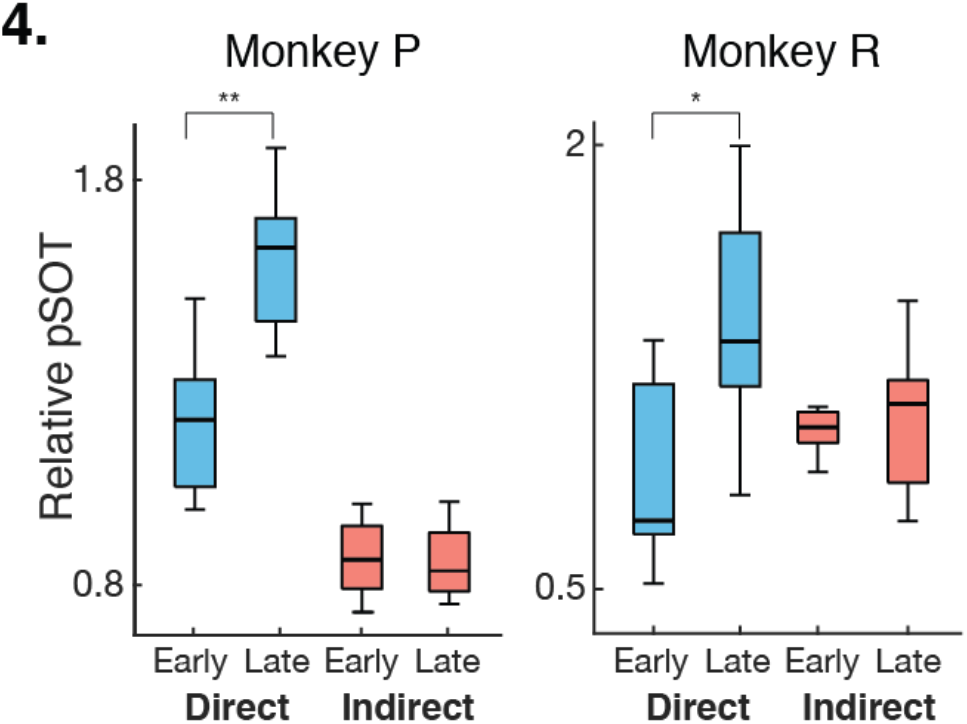
Consolidation is driven by the direct population. Respective contributions of direct and indirect sub-populations to the SOT ratio (pSOT, see Methods for details) in early and late learning relative to contributions in early learning (Unpaired t-test; Monkey P, direct p = 0.002, indirect p = 0.857; Monkey R, direct p = 0.022, indirect p = 0.624).

To characterize how neural exploration modified the direct and indirect neural subspaces differently, we quantified these changes by calculating the shared alignment pairwise between each training epoch’s shared covariance matrix for each subpopulation (Figure 5a). The shared alignment compares how much of the shared space of one epoch projects onto the shared space of another epoch. Intuitively, given two 2-dimensional shared subspaces, the shared alignment compares the angle between the two planes. Likewise, orthogonal planes would result in a shared alignment of 0 and perfectly aligned planes would result in a shared alignment of 1. If the shared subspace consolidates with learning, as has been shown in direct subpopulations (Athalye et al., 2017), we would expect the shared subspace to rotate from the initial subspace over learning. If the shared subspace remains fixed over learning, we would predict that the alignment between the first epoch and later epochs remains high, indicating little change in the coordinated activity of the population. Since we are interested in how the subspaces pertaining to specific populations change over time, we analyzed only the neurons that were stable across learning. We found that the shared alignment decreased from the first epoch for both subpopulations (Figure 5b). This indicates that both subpopulations rotated their low-dimensional subspaces, suggesting that neurons may adapt on a network level that includes both direct and indirect neurons.

**Figure 5.**
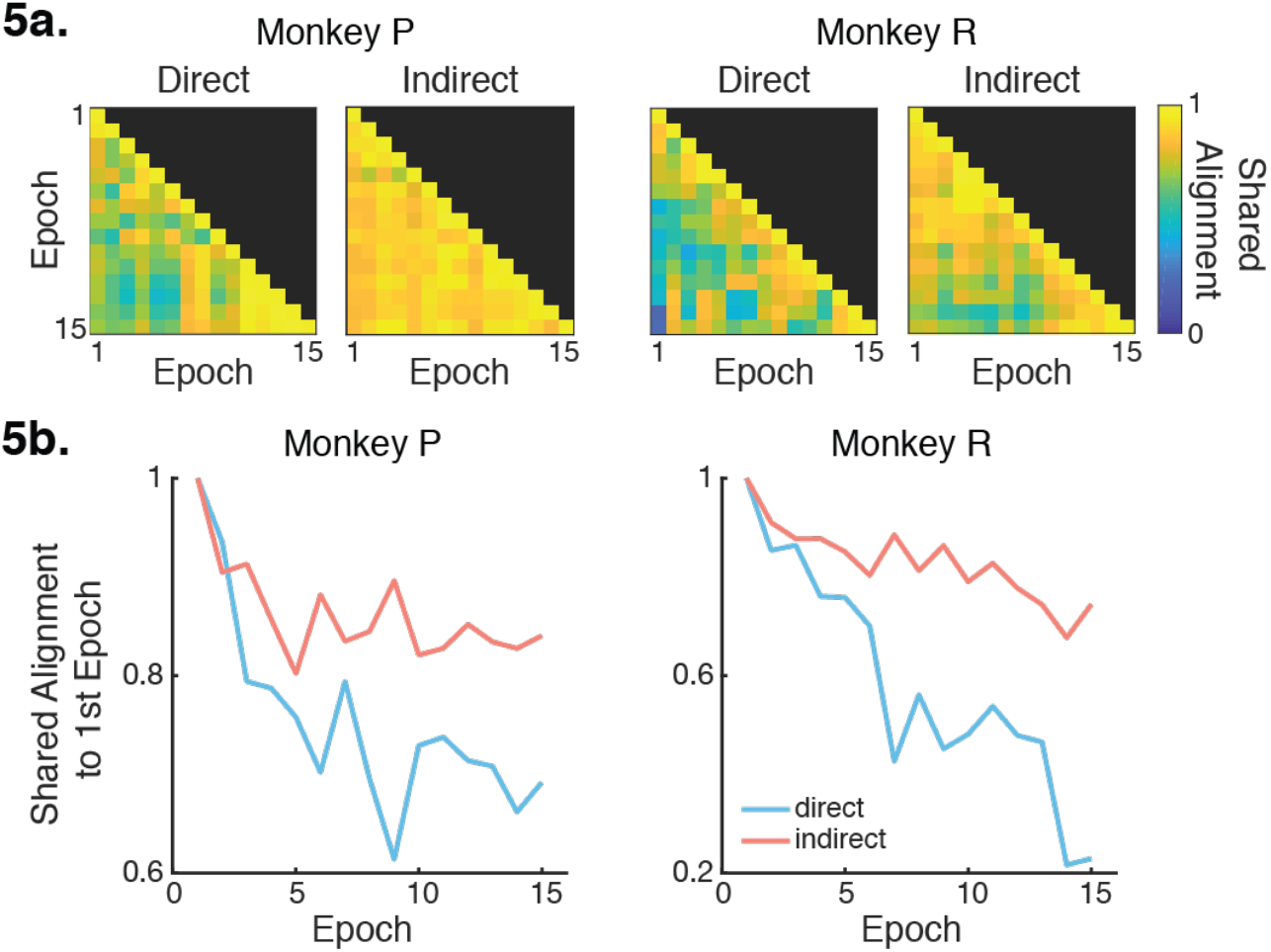
Neurons rotate onto a stable low-dimensional solution subspace. (a) Shared alignment was calculated pairwise between epochs for each subpopulation. (b) Alignment diverges from the first epoch in both subpopulations (Linear regression; Monkey P: direct, R^2^ = 0.530, p = 0.001, indirect, R^2^ = 0.352, p = 0.012; Monkey R: Direct R^2^ = 0.857, p = 4.63e-7, Indirect R^2^ = 0.760, p = 1.41e-5).

To more closely observe how indirect neural activity follows direct neural activity, we analyzed data from a second experiment in which Monkey P learned to perform the same BMI task with a new decoder following proficient control with the original learned decoder (Figure 6a). Eight experimental blocks were performed over the course of four days, alternating between control with the new and previously learned decoder each day. The neural variance for direct and indirect neurons was calculated for each of these eight blocks. Note that only stable indirect neurons were used for this analysis since we wanted to explicitly track how the variance changed as a function of block number. We found that both subpopulations increased and decreased neural variance together over blocks, with similar changes in variance between blocks occurring in both direct and indirect neurons (Figure 6b,c). Thus, improved performance with the new decoder involved increased exploration not only by the direct neurons, but also by the supporting indirect neurons.

**Figure 6.**
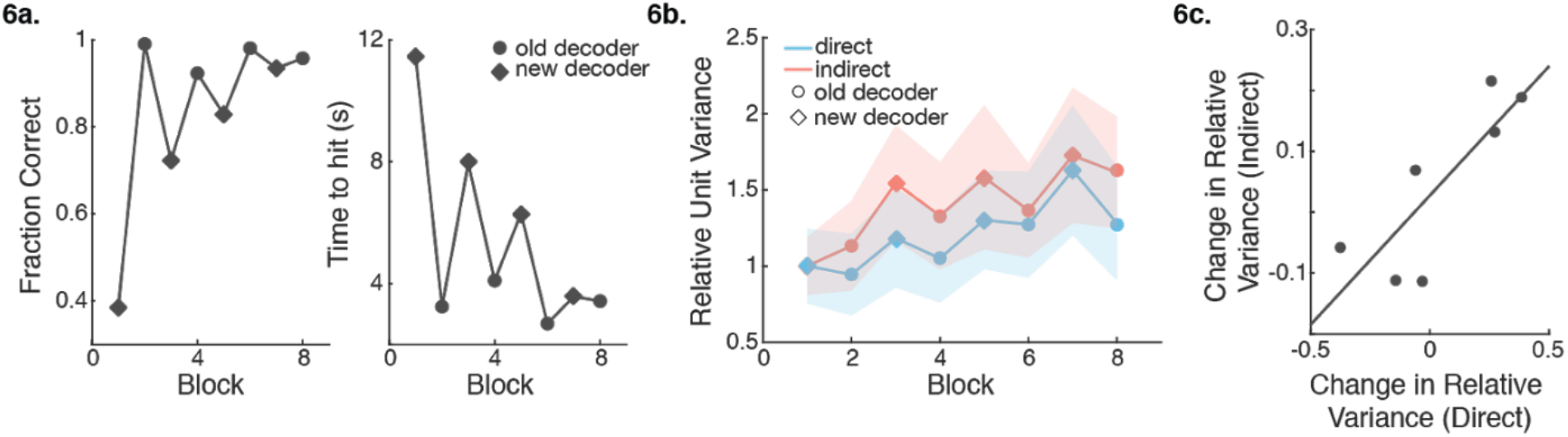
Neural variance modulates concomitantly between subpopulations. (a) Monkey P learns to perform BMI with a new decoder following proficient control with the old decoder. 8 experimental blocks were performed, alternating between a new decoder (diamond) and the previously learned decoder (circle). Fraction of initiated trials that were successful increased over training blocks (left) and the time to reach a target decreased over training blocks (right). (b) Both subpopulations increase their neural variance over blocks (Linear regression; Direct R^2^ = 0.623, p = 0.020, Indirect R^2^ = 0.655, p = 0.015). The relative variances across blocks are correlated between subpopulations (Pearson’s r, r = 0.856, p = 0.007). (c) Each point represents the change in relative variance between two consecutive blocks. The changes in relative variance within the direct and indirect subpopulations are correlated (Pearson’s r, r = 0.820, p = 0.024).

One explanation for why indirect neurons would increase variance alongside direct neurons could be due to physiological projections from other brain regions into M1. It is unlikely that the brain would be able to selectively increase neural variance for only the direct neurons and is much more likely to increase the variance in broader networks involving indirect neurons. To gather an intuition of how broad these networks may be, we separated the indirect neurons recorded during the original 15 epochs in Monkey P into “far” and “near” indirect neurons. “Far” indirect neurons (N = 29-69) were those recorded on electrodes not containing direct neurons. In contrast, “near” indirect neurons (N = 7-14) were indirect neurons that existed on the same electrode shanks as direct neurons. Monkey R was excluded from these analyses due to recording too few near indirect neurons during several epochs (N = 0-10). We found that neural variance increased for both far and near indirect neurons between early and late learning (Figure 7a). However, the pSOT only increased for the near indirect subpopulation and significantly decreased for the far indirect subpopulation (Figure 7b). Together, these results suggest that while neural exploration exists in broader networks consisting of both direct and indirect neurons, neurons closer in proximity to direct neurons consolidate more than those farther away from direct neurons.

**Figure 7.**
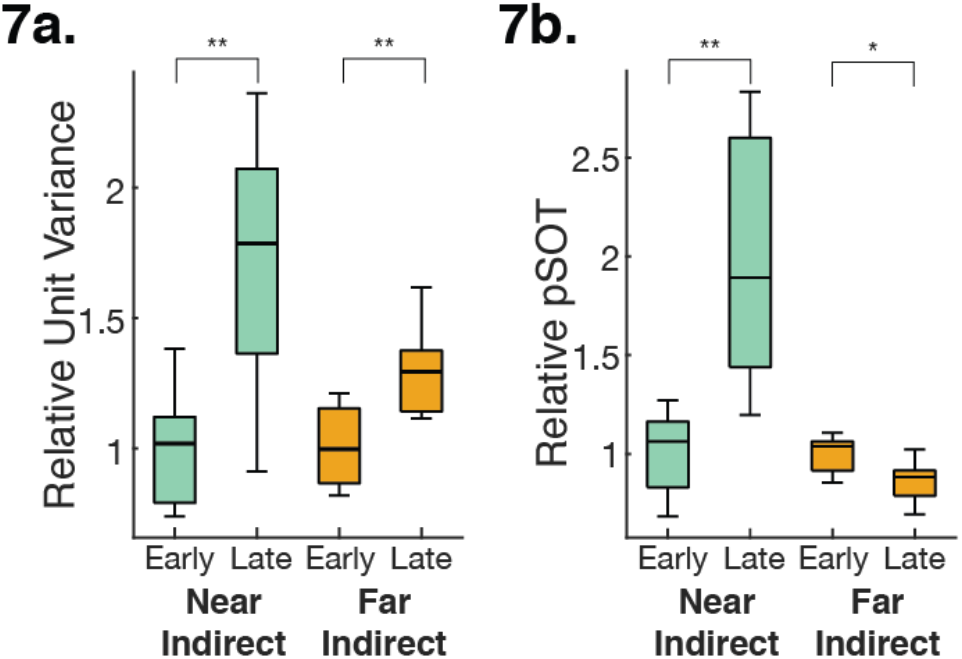
Neurons differentially consolidate based on proximity to direct neurons. (a) Both near and far indirect neurons exhibited significant increases in neural variance (Unpaired t-test; near p = 0.006, far p = 0.007). (b) Only near indirect neurons exhibited a significant increase in pSOT relative to early learning (Unpaired t-test; p = 0.003). Far indirect neurons exhibited a significant decrease in pSOT (Unpaired t-test; p = 0.022).

## Discussion

In this study, we explored how changes in cortical dynamics underlie skill refinement. Using a BMI allowed us to examine how cortical dynamics of subpopulations of neurons change over learning when the mapping of neural activity to behavior is precisely defined. That is, how do both cortical neurons that are used as direct inputs into a decoder (direct neurons) and neurons in the supporting network (indirect neurons) adapt over learning?

Our results showed a concerted effort in exploration over learning where both direct and indirect neurons increased variance in their firing rates over learning (Figure 2). Furthermore, we found that the increased coordination of neural firing patterns existed in the combined population (Figure 3). Previous work has suggested that these changes are characteristic of more stereotyped behavior over learning and this increase in coordination in the population has been shown to lead to more direct paths to target (Athalye et al., 2017). Thus, when considering the entire recorded population, it appears as though the combined population adapts together to facilitate learning. However, past work has shown differences in adaptation between direct and indirect neurons (Ganguly et al., 2011; Gulati et al., 2014; Koralek et al., 2013). When we considered the relative contribution of each subpopulation to the increase in coordination, we found that the indirect subpopulation contributed very little to the overall changes in coordinated patterns. (Figure 4). That is, while we witnessed an increase in coordination in the entire recorded population, there was less within-group coordination in indirect neurons compared to direct neurons. Our metrics of coordination (e.g., SOT) rely on averaging the amount of correlated activity between pairs of neurons. Consolidation occurring more heavily in one subpopulation would nevertheless increase the SOT in the entire population. Altogether, while both subpopulations exhibit similar levels of exploration over learning, the exploration by the indirect subpopulation results in less increases in coordination of neural firing patterns than that of the direct subpopulation.

The coordination of neural activity resulted in rotations of the neural space over learning in both direct and indirect subpopulations (Figure 5). These rotations can be intuitively thought of as the correlated co-firing patterns of neurons and rotating the neural space corresponds to adjusting which neurons are more active given the state of other neurons in the population. Following this interpretation, both direct and indirect subpopulations similarly adapted their coordinated firing patterns within subpopulation over learning. Furthermore, when we examined how both subpopulations changed in an experiment where two decoders were swapped each day, we found that changes in neural firing rate variance were proportional between the two subpopulations (Figure 6). Increases in neural variance in the direct subpopulation were mirrored by similar changes in the indirect subpopulation when switching to the less familiar decoder. This suggests that neurons within the supporting cortical network may be adapting in similar ways to the direct neurons. Consequently, there may not be a clear distinction between direct and indirect neural activity, but rather the two subpopulations are adapting via the same mechanism to different extents. However, this experiment was limited to only four days of switching between decoders. It is unclear whether or not these parallel changes in neural variance between the direct and indirect subpopulations would continue if the animal was given more extensive practice.

Our findings provide further evidence for existing hypotheses on how the brain learns to refine coordinated neural dynamics (Athalye et al., 2020). Specifically, small networks of cortical neurons may be driven by upstream subcortical inputs. We found that indirect neurons adapted in similar ways as direct neurons, suggesting that some indirect neurons may in fact be adapting with or alongside direct neurons. When we asked how distance from direct neurons influenced these results, we found that indirect neurons in closer spatial proximity to direct neurons increased coordination more than indirect neurons that were farther away (Figure 7b). While the upstream projections from subcortical structures are not necessarily spatially organized, our results are consistent with the hypothesis that subcortical structures may be driving changes in smaller groups of neurons (Athalye et al., 2020). Furthermore, it has previously been shown that when disparities are present between the control space and neural space (i.e. how well the decoders aligned to the natural firing patterns of the neurons), neurons with larger disparities adapt more over learning compared to neurons with smaller disparities (Athalye et al., 2017; Chase et al., 2012; Jarosiewicz et al., 2008; Orsborn et al., 2014). How might the nervous system explore and refine coordinated population dynamics across a period of multiple days? A recent study has shown that task-related neurons, consisting of direct neurons as well as task-modulated indirect neurons, increase coherency to slow-wave activity (SWA) during sleep which has been linked with consolidation (Gulati et al., 2014). This suggests that in addition to online task practice, neural reactivations during sleep can aid in exploring the contributions of direct and indirect neural population relative to successful outcomes and reward. Notably, indirect neurons that were closely tied to reward were preserved and resembled direct neurons; this might explain why some indirect neurons were modified during neuroprosthetic skill acquisition. This also provides further evidence that mechanisms of reinforcement learning may underlie our observed phenomena. Thus, it is quite plausible that adaptation of neural activity over BMI learning is attributed to finding the clusters of neurons with a direct effect on behavior, which may include both direct and indirect neurons, depending on their specific network connectivity and temporal association with successful outcomes.

In this study, we used factor analysis to find correlations in the neural activity of the recorded population. Underlying this model are latent factors, variables that coarsely group neurons together based on coordinated activity patterns. Importantly, the activity of a single neuron can be associated with multiple latent factors. While we were agnostic to what the latent factors in FA may correspond, they may be physiologically analogous to upstream connections from subcortical structures that drive changes in small clusters of neurons that contain direct neurons.

The idea that neural reinforcement is dependent on cortico-striatal circuits, similar to behavioral reinforcement, has been previously supported by studies using BMIs. For example, as rodents learned to produce specific patterns of cortical activity, coherence between these neurons and dorsal striatum emerged and neurons in dorsal striatum developed target-predictive modulation of firing activity (Koralek et al., 2012, 2013; Neely et al., 2018). Furthermore, mice without functional NMDA receptors in striatal projection neurons could not learn to re-enter a cortical pattern that led to reward. Thus, cortico-striatal plasticity is necessary for learning to efficiently produce the cortical activity patterns required to obtain rewards. These findings along with the results from our study further support the hypothesis that smaller clusters of neurons, which may include both direct and indirect neurons, are adapted over learning more than clusters of neurons that do not drive behavior. Future work involving simultaneous recordings from more cortical and subcortical areas as well as experiments recording from larger populations of neurons are imperative to understand how this selection process occurs.

Overall, our results demonstrate that the brain learns to modify cortical population dynamics in subpopulations relevant for behavioral control. When using a BMI, we find that neurons with direct input to the decoder as well as neurons in the surrounding cortical network increase exploration and consolidate onto a low-dimensional neural space. The degree of this consolidation is dependent on the relationship of the neural activity to the behavioral output. Thus, the brain may not be reinforcing the activity of single neurons, but rather reinforcing cortical population dynamics that are relevant to producing a desired behavior. These findings indicate that the brain learns to control a BMI by refining cortical population-level dynamics, suggesting that BMI decoders extracting information based on population-level statistics, such as the covariance structure of the population, may be more effective compared to traditional decoding methods based on the statistics of individual neurons. Understanding the role of modifications of adjacent indirect activity in obtaining precise control of a BMI may help us understand the neural adaptation that is required for achieving long-term, stable control of a BMI.

## Methods

### Animal Subjects

All procedures were conducted in compliance with the NIH Guide for the Care and Use of Laboratory Animals and were approved by the University of California at Berkeley Institutional Animal Care and Use Committee.

Two adult male rhesus monkeys (Macaca mulatta) were chronically implanted in the brain with arrays of 64 microelectrodes (Innovative Neurophysiology, Durham NC) (Ganguly & Carmena, 2009). Monkey P was implanted in the left hemisphere in the arm area of both primary motor cortex (M1) and dorsal premotor cortex (PMd), and in the right hemisphere in the arm area of M1, with a total of 192 microwires across three implants. Monkey R was implanted bilaterally in the arm area of M1 and PMd (256 microwires across four implants). Only activity from M1 was included in the direct ensembles (Monkey P: right M1; Monkey R: left M1) and only activity from the same hemisphere was included in the indirect ensemble. Array implants were targeted for pyramidal tract neurons in layer 5. Localization of target areas was performed using stereotactic coordinates from a neuroanatomical atlas of the rhesus brain (Paxinos et al., 2000).

### Electrophysiology

Neural activity was recorded using the MAP system (Plexon, Dallas TX). Stable units, to be part of the direct ensemble, were selected based on waveform shape, amplitude, relationship to other units on the same channel, interspike interval distribution, and the presence of an absolute refractory period. Only units from primary motor cortex were used which had a clearly identified waveform with signal-to-noise ratio of at least 4:1. Activity was sorted prior to recording sessions using an online spike-sorting application (Sort Client; Plexon). Stability of waveforms was confirmed by analyzing the stability of PCA projections over days (Wavetracker; Plexon).

Direct units are defined as the units being used to control the BMI. Indirect units consisted of the remaining recorded units. For analyses including only stable units from the same hemisphere, stability in the indirect ensemble was assessed using pairwise cross-correlograms, autocorrelograms, waveform shapes, and mean firing rates (Fraser & Schwartz, 2012).

### Experimental Setup and Behavioral Training

#### Manual Control Training Before BMI

Before starting the BMI learning experiments, subjects were overtrained on the task performed with arm movements using a Kinarm (BKIN Technologies, Kingston ON) exoskeleton which restricted shoulder and elbow movements to the horizontal plane.

#### BMI Tasks

Data from Ganguly & Carmena, 2009, in which subjects performed a self-initiated, eight-target, center-out reaching task, was analyzed. In these experiments, a cursor on a screen was continuously controlled by neural activity. Subjects self-initiated trials by moving the cursor to a center target. One of the eight peripheral targets was randomly selected each trial. Self-initiated trials consisted of those in which the animal moved the cursor to the center target and held for 250-300ms. Successful trials required the animal to move the cursor to the peripheral target within 15s of initiating the trial and hold the cursor at the target for 250-300ms. Successful trials resulted in a juice reward; failed trials were repeated. During BMI control, both arms were removed and lightly restrained. Neither animal moved their upper limbs during BMI control.

After the initial 19-days of performing the BMI task with a fixed-decoder, Monkey P learned a second decoder over the course of four days. Within each of these four days, Monkey P performed one training block of the new decoder, followed by one training block of the old decoder. Both of these decoders were fixed and used the same direct ensemble as input.

### Preprocessing Pipeline

For all analyses, neural data was binned into 100ms bins to match the decoder timescale. Additionally, learning was analyzed over “training epochs,” where each epoch consisted of 150 self-initiated trials. Learning took place over the first 15 training epochs (2250 self-initiated trials). We chose to analyze the data across training epochs, rather than days, to eliminate the effect of variable numbers of trials each day. Only the first 15 training epochs (2250 self-initiated trials) were analyzed; we defined early and late learning as the first and last seven of these training epochs, respectively. Monkey P initiated a total of 3589 trials. Monkey R initiated a total of 2357 trials.

### Factor Analysis

#### Shared-Over-Total Variance Ratio

Factor analysis (FA) was conducted on the neural population for each epoch to observe underlying correlated neural activity. FA decomposes population signals into correlated and uncorrelated components. For a given neuron *i*, correlated activity is represented by the shared variance 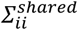, and the degree to which the activity was correlated over learning was represented by the ratio of shared-over-total variance of the neural population (SOT). We calculated the SOT ratio according to the methods described in Althaye et al., 2017.

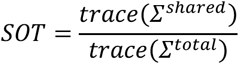

#### Total Variance

We also considered how the total variance changed between early and late learning. This was the sum of the private and shared variances 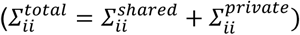.

#### Partial Shared-Over-Total Variance Ratio

We quantified the respective contributions of subpopulations to the SOT ratio using the partial shared-over-total variance (pSOT) ratio. Here, we compared the sum of the shared variance for each subpopulation over the total variance for the entire population. A relative measure was used to account for the fact that the direct and indirect ensembles were different sizes. That is,

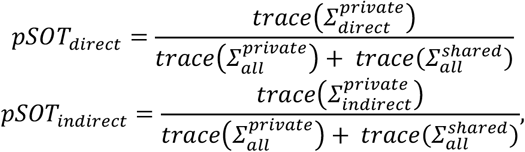

where *Σ_i_* denotes the covariance matrix for the (sub)population *i*.

#### Shared Space Alignment

We used the ‘‘shared space alignment’’ to measure the similarity between the shared variance of two different training epochs. The shared space alignment is the fraction of shared variance from one epoch captured in the shared space of a second epoch and thus ranges from 0 to 1. We calculated shared alignment according to the methods described in Althaye et al., 2017. Given two epochs, A and B, we first compute the projection matrix into Epoch B’s shared space, *col*(*U^B^*). We then project *Σ^A,shared^* onto B’s shared space, 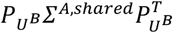. Finally, the alignment is calculated,

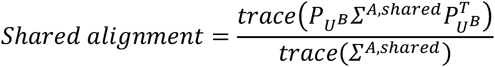

#### Quantification & Statistics Analyses

All analyses were performed on each training epoch separately. Trends were analyzed for significance with linear regressions. Epochs 1-7 and Epochs 9-15 were grouped into “early” and “late,” respectively. Epoch 8 was omitted so that there were equal numbers of epochs for both early and late learning. Groupings of early and late epochs were compared using an unpaired two-sample t-test.

## Acknowledgments

We thank N. Vendrell-Llopis for helpful comments on the manuscript and V. Athalye for valuable discussions. This work was supported by the National Science Foundation Graduate Research Fellowship (to E.L.Z. and to A.K.Y.), the Department of Veterans Affairs, Veterans Health Administration, Rehabilitation Research and Development, and the American Heart Association/American Stroke Association (to K.G.), the Alfred P. Sloan Foundation, the Christopher and Dana Reeve Foundation, the National Science Foundation CAREER Award #0954243, the Defense Advanced Research Projects Agency contract N66001-10-C-2008, and the National Institute of Health Award R01NS106094 (to J.M.C.).

